# Micromorphy Offers Effective Defence Against Predation: Insights From The Cost-Benefit Analyses Of Micro Gastropod Predation Record

**DOI:** 10.1101/2021.05.30.446364

**Authors:** Anupama Chandroth, Devapriya Chattopadhyay

## Abstract

Predation, an important driver of natural selection, is studied in the fossil record using quantifiable traces like drill holes produced by gastropods and repair scars produced after durophagous attacks. Despite the abundance of such records in molluscan prey, predation records of micromolluscs (<5mm) remained unexplored. Using a Miocene assemblage of microgastropods from the Quilon Limestone, India, we established the predatory-prey dynamics with the help of cost-benefit analyses. The overall predation intensity is low (DF = 0.06, RF= 0.04) and does not depend on the relative abundance of prey groups suggesting a non-random prey selection regardless of the encounter frequency. The predation is selective in terms of taxonomy, ornamentation, and size of the prey. The smallest size class has the lowest DF and RF supporting a negative size refugia. Higher IDF in larger size class and ornamented groups implies morphological defense resulting in higher failure. Microgastropods show a lower predation intensity than their regular-sized counterparts in a global comparison of coeval records. Results of the cost-benefit analyses explain this difference; the net energy gain from predatory drilling is found to increase monotonically with increasing prey size making the small prey less beneficial. Because the predators try to maximize net energy gain from a predatory attack, the microgastropod prey characterized by relatively low net energy yield is not preferred in the presence of larger prey. Micromorphy, therefore, appears a viable strategy for the prey group to be adopted as an evolutionary response against predation, especially in resource-limited conditions that fail to support large body size.

## INTRODUCTION

Predation is an important ecological interaction and one of the major drivers of natural selection (Kelley and Hansen, 1993; Kitchell, 1986; Vermeij, 1987). It also plays a vital role in shaping the community structure (Barnes et al., 2010; Hines et al., 1990). However, it is challenging to find traces of such interactions in the fossil record that can be studied quantitatively. Trace fossils like predatory drill holes and repair scars are common evidence of predation in fossil records (Kelley et al, 2003). Complete drill holes represent a lethal attack in contrast to the traces of non-lethal attacks such as incomplete drill holes, repair scars. These traces reveal various aspects of predation (including the predatory identity, prey preference, success rate) (Klompmaker et al., 2019). The fossil record of predatory traces proved crucial in understanding the evolution of marine invertebrates and restructuring of the marine ecosystem as a response to biotic interaction (Vermeij et al. 1981; Huntley and Kowalewski, 2007).

The relative size of the prey and its predator often determines the outcome of a predatory interaction and plays an important role in shaping the evolutionary trajectory of the groups (Klompmaker et al, 2017; Vermeij 1987). In drilling predation, the prey size preference is primarily governed by the energy maximization of the predator for each attack (Chattopadhyay and Baumiller, 2009; Kitchell et al, 1981). Patterns like size refugia are common among the invertebrates where prey beyond a specific size class are seldom attacked. Predators avoid preying these large prey species because the capture involves the investment of more energy than the gain rendering the predation nonbeneficial (Harper et al, 2009; Leighton, 2002). Small prey is not always the most preferred size class either. Smaller brachiopods demonstrated the lower intensity of shell-breaking predation (Harper et al, 2009). A fossilized assemblage of micro bivalves also revealed a lower intensity of drilling predation in the smaller size class supporting the existence of a negative size refugia (Chattopadhyay et al, 2020). Such predation resistance among extremely small invertebrates points to a complex relationship between size and predation intensity. To understand the evolutionary response of small size to predation, the predation record of micromolluscs needs to be explored. Except for the microbivalves, the predation record of microfossil primarily constitutes of taxa such as foraminifera (Culver and Lipps 2003), ostracods (Maddocks 1988; Rayment and Elewa 2003); microgastropods have not been studied for their predation record.

Here we studied the microgastropods from the early Miocene seagrass bed from southwest India (Quilon, Kerala) (Harzhauser 2014) to address the following questions:

1. What controls the prey selectivity in microgastropods?
2. Does the predation in microgastropod viable from the cost: benefit perspective?
3. Is the predator-prey dynamics significantly different in microgastropods in comparison to the macrogastropods?

## MATERIALS AND METHODS

### Locality and collection

The field locality is situated on the cliffs along the shores of Ashtamudi lake, near Padapakkara village, Kerala, India (N 98 08° 58’36”, E 076° 38’08”) (Figure 1). The collection protocol has been described in detail in Chattopadhyay et al (2020).

**FIGURE 1.**
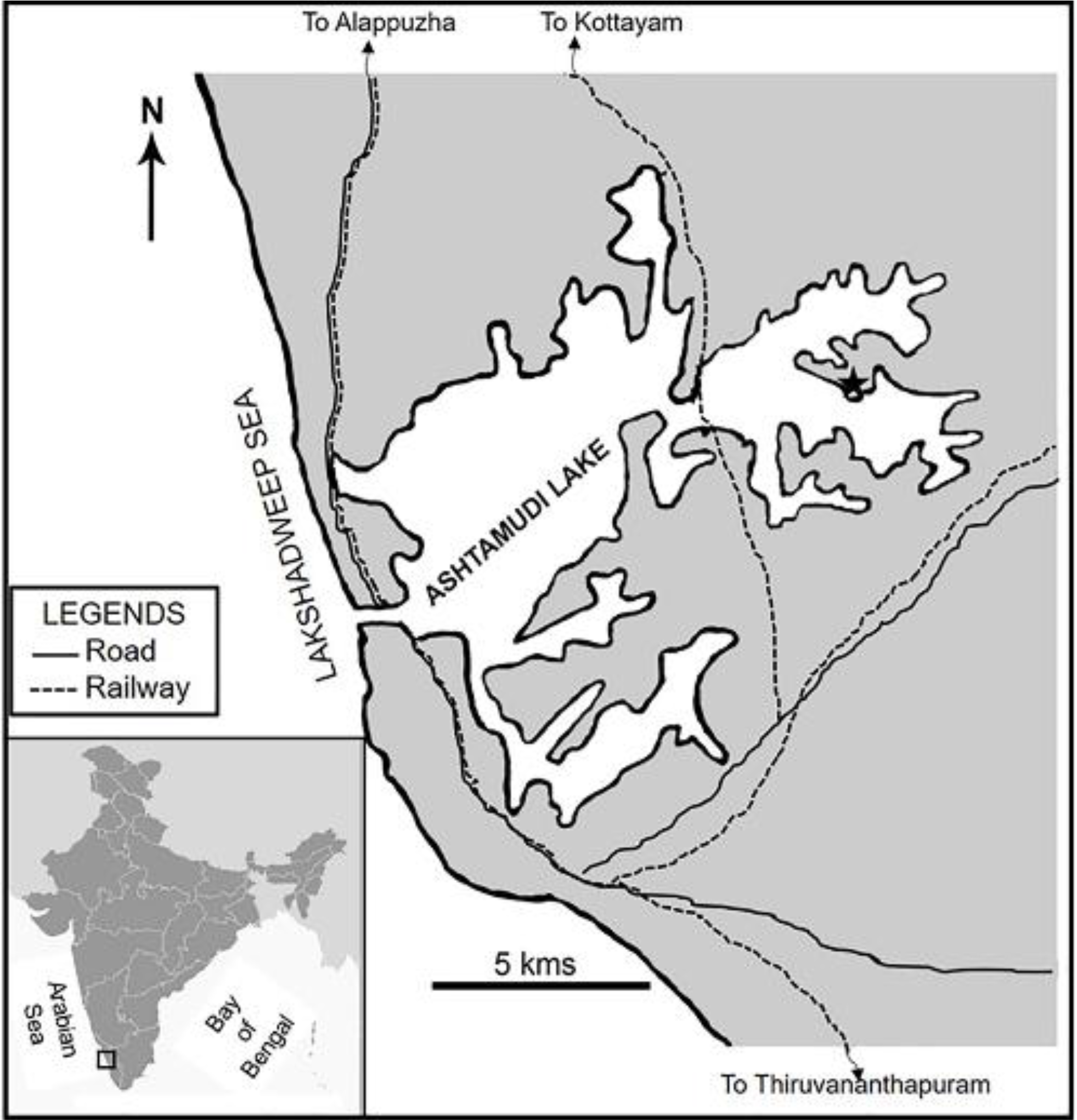
Location of the studied locality with the map of India (inset). The star represents the location of the Quilon Limestone bed (Kerala) (modified after Chattopadhyay et al, 2020).

From the collected bulk sample, 371.8g of the sample was processed. The bulk sample was soaked in normal water for 5-6 days to loosen the sediments and subsequently, wet sieved using an 18μm sieve to remove the sedimentary particles. The remaining sediments were then dried, sieved, and classified into different size classes using a set of 5 sieves (mesh sizes 63, 60, 35, 25, 18μm). We studied the processed samples under the microscope and identified them up to the family level using the detailed study by Dey (1961) and Harzhauser (2014). The identified specimens were categorized into three size classes, small (less than 1mm), medium (1-2mm), and large (greater than 2mm). The specimens were also classified into two groups based on ornamentation: the ones with smooth shells were classified as non-ornamented (Buccinidae, Eulimidae, Marginellidae, Naticidae, Phasianellidae, Scaliolidae, and Turbinidae) and rest as ornamented. We used two protocols for characterizing the location of drill holes. In the first protocol, the gastropod shell is divided into two equal zones radially (apertural and abapertural) and each drill hole site is characterized using this scheme. In the second protocol, the gastropod shell is divided into three sections vertically (top, central, basal) from the apex. Considering the total height of a specimen, three sections are assigned based on the relative distance from the apex as the top (33% at the top), basal (33% at the base), and central (remaining 33% at the center). We took detailed photographs of drilled specimens using a Nikon D700 attached with an Olympus SZX16 microscope. We process the images using Image J to measure the size of the specimens and drill holes. Drilling predation on Miocene macrogastropods from the same biogeographic region (Goswami et al, 2020) and other localities (Hoffmeister and Kowalewski, 2001; Kelley and Hansen, 2006; Sawyer and Zuschin, 2011) were compiled for comparative analysis.

### Analysis

Drilling frequency (DF), a measure of successful attempt, is calculated by dividing the number of drilled specimens by the total number of specimens.

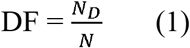

Where, N_D_ = number of specimens with complete drill hole

N= Total number of specimens.

The incomplete drilling frequency (IDF) is calculated by dividing the total number of incomplete drill holes by the total number of drilling attempts (Chattopadhyay & Dutta, 2013).

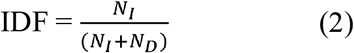

Where, N_D_ = number of specimens with complete drill hole

*N_I_* = number of incomplete drill holes

To estimate the intensity of repair scar (RF), the total number of specimens with repair scar was divided by the total number of individuals.

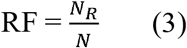

Where, N_R_ = number of specimens with repair scar

N= Total number of specimens.

To estimate the occurrence of multiple predation traces, we calculated MULT as the total number of holes in the specimens with multiple drillholes, divided by the total number of drilling attempts (Kelley and Hansen, 1993).

We used the Pearson correlation test to evaluate the correlation of predation intensity with abundance and size. We used a two-tailed chi-square test to evaluate the variation in predation intensity (DF, IDF, and RF) between different size classes. For the site preference of drilling a chi-square test of goodness of fit was done. All the statistical tests are conducted using the R programming environment (R development core team 2007).

### Cost-benefit analyses

We reconstructed the size of the predator (*Lpd*) from the drillhole size (*OBD*) using the following equation proposed by Klompmaker et al (2017).

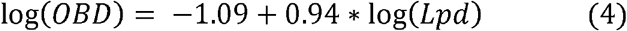

*OBD* = *Outer borehole diameter (mm)*

*Lpd* = *Length of the gastopod predator* (*mm*)

The cost-benefit analysis was done for the microgastropods by adapting the equation suggested by Kitchell et al (1981), along with a few modifications. The total benefit is calculated using the ash-free dry weight (*Wpr*) of gastropod prey with a specific size (*Lpr*). We used the formula for the genus *Polinices* for all the species. The relation is given as (Edwards and Huebner, 1977)

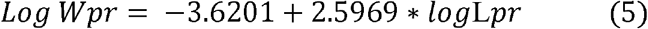

Where

*Wpr* = *Ash free dryw eight of the prey* (*g*)

*Lpr* = *Length oft he gastopod prey* (*mm*)

The calculated ash-free dry weight (Eqn.5) is then multiplied by the energetic conversion factor, 21.46kj/g (Kitchell et al 1981) to obtain the benefit.

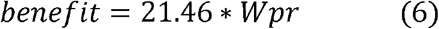

The cost is calculated as a product of metabolic rate and time taken to drill the prey species. The drilling time (t) is found to be directly related to the thickness of the shell (T). The shell thickness (T) is calculated as (Avery and Etter, 2006)

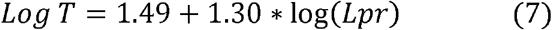

Where

*T* = *Thickness of the shell* (*μm*)

*Lpr* = *Length of the gastropod prey* (*mm*)

Using the thickness (T), we calculated the time (t) required to produce the drill hole (Kitchell et al. 1981)

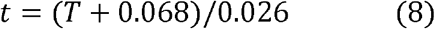

Where

*T* = *Thickness* (*mm*)

*t* = *drilling time* (*hours*)

The metabolic rate of the predator is estimated through a series of steps. Using the OBD, the length of the predator (*Lpd*) is calculated (Eqn. 4).

Later the ash-free dry weight is calculated using the following relationship:

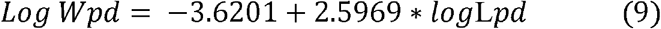

Where

*Wpd* = *Ash free dryw eight of the predator* (*g*)

*Lpd* = *Length oft he gastopod predator* (*mm*)

The ash-free dry weight (Eqn. 9) is then used to find the metabolic rate in terms of the amount of oxygen consumed per hour (Harper and Peck 2003)

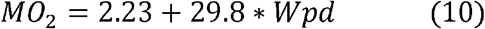

Where

*MO*_2_ = *Amount of oxygen consumed* (*μg*)

*Wpd* = *Ash free dry weight of the predator* (*g*)

According to Harper and Peck (2003), 18.6μg of oxygen/ hour is equivalent to 13μl of oxygen/ hour. This relation is used to calculate the amount of oxygen consumed in litres. Using standard conversion factors, we obtain the metabolic rate in kJ/hour.

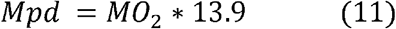

Where

*Mpd* = *Metabolic rate of predator* (KJ/hour)

The cost is estimated as

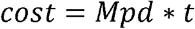

Using eqn. 6, the net energy gain is estimated from the following expression:

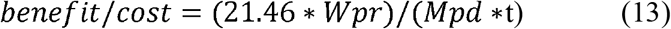

## RESULTS

Out of a total of 1328 microgastropod specimens in our study representing 39 species, 35 genera, and 25 families. Cerithiidae is the most abundant family, represented by 718 individuals, followed by Pyramidellidae and Scaliolidae. A total of 150 individuals from 14 families show the signature of predation yielding an overall DF of 0.063 and RF of 0.039 (Table 1, Figure 2, 3). Eleven families, represented by more than ten individuals each, are considered for subsequent predation analyses (Figure 2, 3, Table1). Multiple drill holes are found on 5 specimens representing 3 different families, namely Rissoinidae, Cerithidae, and Pyramidellidae. The MULT of the overall assemblage is 0.0971.

**TABLE 1.** Overall abundance and summery of drilling predation of micrograstropods from Quilon Limestone bed.

| Family | Total specimens | Drilled specimens | Drilling frequency (DF) | Specimens with incomplete drilling (ID) | Incomplete drilling frequency (IDF) | Specimens with repair scars | Repair frequency |
| --- | --- | --- | --- | --- | --- | --- | --- |
| Cerithiidae | 717 | 34 | 0.047 | 4 | 0.105 | 13 | 0.018 |
| Pyramidellidae | 195 | 12 | 0.061 | 2 | 0.143 | 21 | 0.108 |
| Scaliolidae | 117 | 1 | 0.008 | 0 | 0 | 2 | 0.017 |
| Rissoinidae | 103 | 23 | 0.223 | 3 | 0.1154 | 3 | 0.029 |
| Eulimidae | 28 | 1 | 0.036 | 1 | 0.5 | 2 | 0.071 |
| Naticidae | 27 | 3 | 0.111 | 0 | 0 | 2 | 0.074 |
| Obtortionidae | 26 | 1 | 0.038 | 3 | 0.75 | 2 | 0.077 |
| Phasianellidae | 23 | 3 | 0.13 | 1 | 0.25 | 2 | 0.087 |
| Turbinidae | 18 | 1 | 0.056 | 0 | 0 | 3 | 0.167 |
| Buccinidae | 13 | 1 | 0.077 | 0 | 0 | 0 | 0 |
| Marginellidae | 11 | 0 | 0 | 0 | 0 | 0 | 0 |
| Olividae | 9 | 2 | 0.222 | 0 | 0 | 0 | 0 |
| Raphitomidae | 9 | 1 | 0.111 | 0 | 0 | 0 | 0 |
| Horaiclavidae | 8 | 1 | 0.125 | 0 | 0 | 1 | 0.125 |
| Triphoridae | 8 | 0 | 0 | 0 | 0 | 0 | 0 |
| Turritellidae | 5 | 0 | 0 | 0 | 0 | 0 | 0 |
| Mangellidae | 2 | 0 | 0 | 0 | 0 | 0 | 0 |
| Columbellidae | 2 | 0 | 0 | 0 | 0 | 1 | 0.5 |
| Borsoniidae | 1 | 0 | 0 | 0 | 0 | 0 | 0 |
| Cerithiopsidae | 1 | 0 | 0 | 0 | 0 | 0 | 0 |
| Epitoniidae | 1 | 0 | 0 | 0 | 0 | 0 | 0 |
| Torchidae | 1 | 0 | 0 | 0 | 0 | 0 | 0 |
| Tornidae | 1 | 0 | 0 | 0 | 0 | 0 | 0 |
| Pseudomelatomidae | 1 | 0 | 0 | 0 | 0 | 0 | 0 |
| Ringiculidae | 1 | 0 | 0 | 0 | 0 | 0 | 0 |
| Total | 1328 | 84 | 0.063 | 14 | 0.143 | 52 | 0.039 |

**FIGURE 2.**
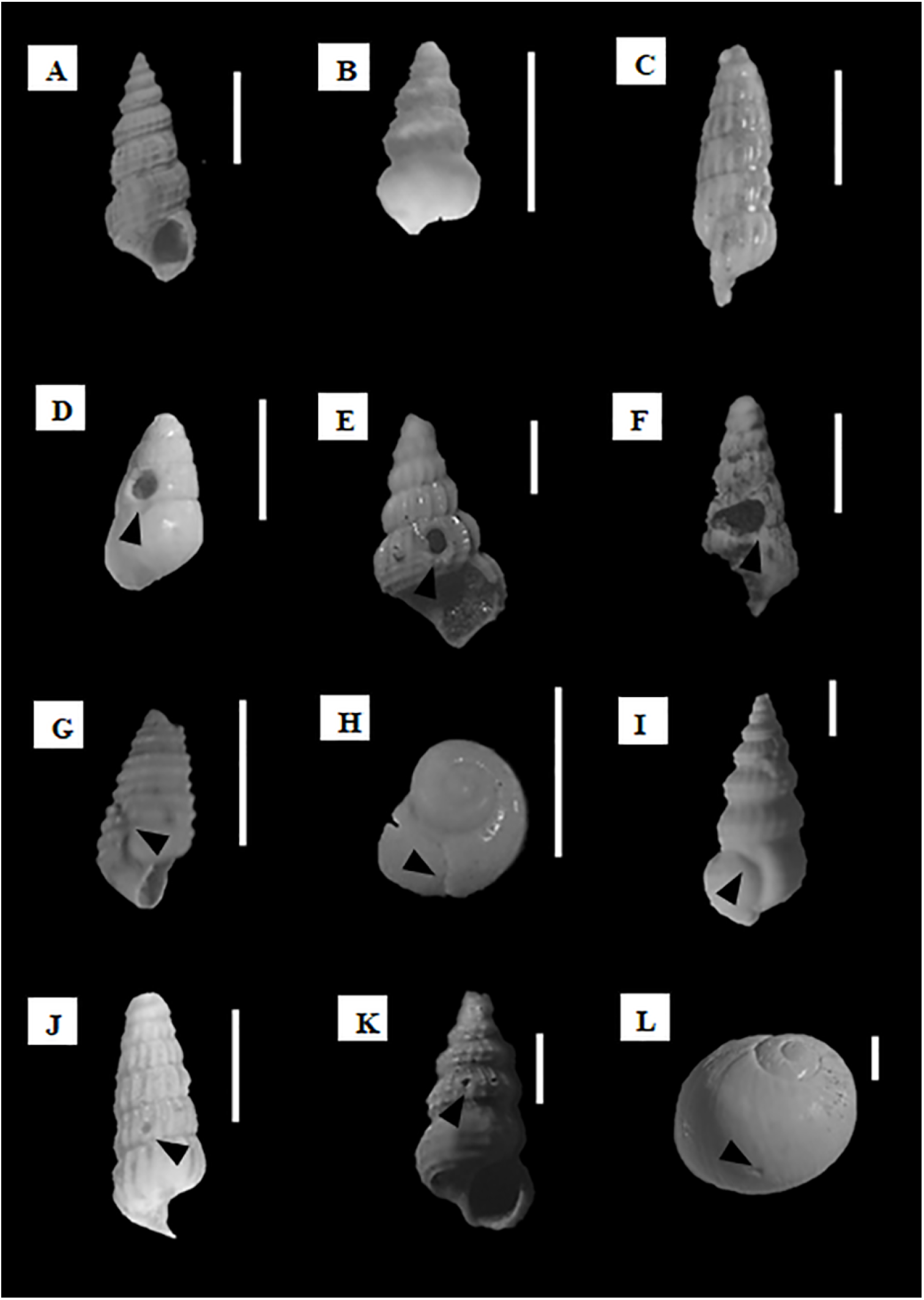
Common gastropod families A) Cerithiidae, B) Scaliolidae and C) Pyramidellidae, gastropods with complete drill hole, D) Phasianellidae, E) Rissoinidae, F) Pyramidellidae; repair marks, G) Pyramidellidae, H) Turbinidae, I) Cerithiidae; incomplete drill hole J) Pyramidellidae, K) Cerithiidae, and predator L) Naticidae. The scale corresponds to 1 mm.

**FIGURE 3.**
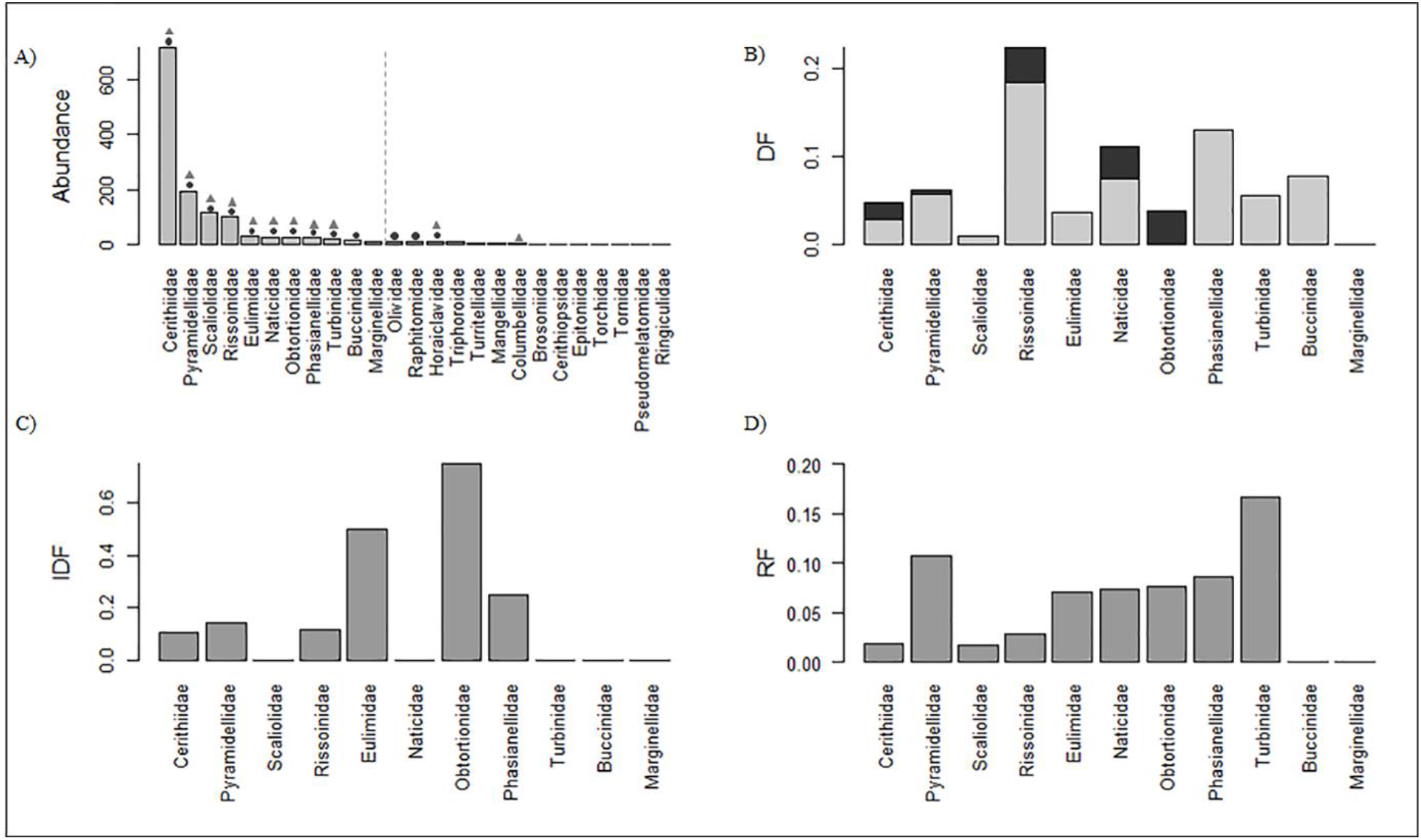
Histograms representing the A) abundance of all families. The dotted line separates the abundant and non-abundant families. The circles and triangles mark those families with drill holes and repair marks, respectively. Histograms representing the B) DF (the darker represents muricids and the lighter naticids), C) IDF and D) RF of the eleven abundant gastropod families.

Among the eleven abundant families, we find ten with complete drilling (Figure 2, Figure 3 B), six with incomplete drill holes (Figure 2, Figure 3C), and nine with repair scars. Rissoinidae and Obtortionidae have the maximum DF (0.22) and IDF (0.75) respectively. The majority of the drill holes correspond to naticid drilling (76.5%) and the rest corresponds to muricid drilling (Figure 3B). Rissoinidae, Cerithiidae, and Pyramidellidae showed multiple drill holes (MULT=0.0971). The overall RF is 0.039 and Turbinidae has the highest RF (0.18). There is no significant correlation between the overall abundance of a family and the observed predation intensity (DF, IDF, and RF) (Figure 4 (A-C)). There is no significant variation in DF or RF between families with and without ornamentation (Figure 4 (D-F)); however, the ornamented shells show a significantly higher IDF (p-value = 0.03) (Figure 4E). The DF, IDF is significantly higher in the larger size (Table 2, Figure 5); RF shows a similar pattern although statistically insignificant (Table 3). The average size of the incompletely drilled specimens is larger than the complete and undrilled specimens (Figure 6A). The apertural placement of complete drill holes is significantly higher compared to the abapertural placement (Chi-square test, p=0.01); apertural placement is least favored for incomplete drilling (Figure 6B). The central region of the shell records the highest incidences of drill holes (79%) (Figure 6C). There is a strong positive correlation between the OBD and the prey size (p < 0.05), especially for naticid predation (p<0.05) (Figure 7 A, B). The overall prey-predator size ratio for microgastropods falls between 0.4 and 1.2 (Figure 7 B). However, ‘small’ microgastropods have a higher prey-predator ratio compared to ‘medium’ and ‘large’ ones (Figure 7). The cost-benefit analysis demonstrates a benefit: cost > 1 for all the successful predation (Figure 7 (C-E)) and this ratio increases with an increase in the size of the prey. The naticid drillings yielded a higher benefit: cost ratio than muricid drilling (Wilcox test, p=0.04) (Figure 7 D, E)

**FIGURE 4.**
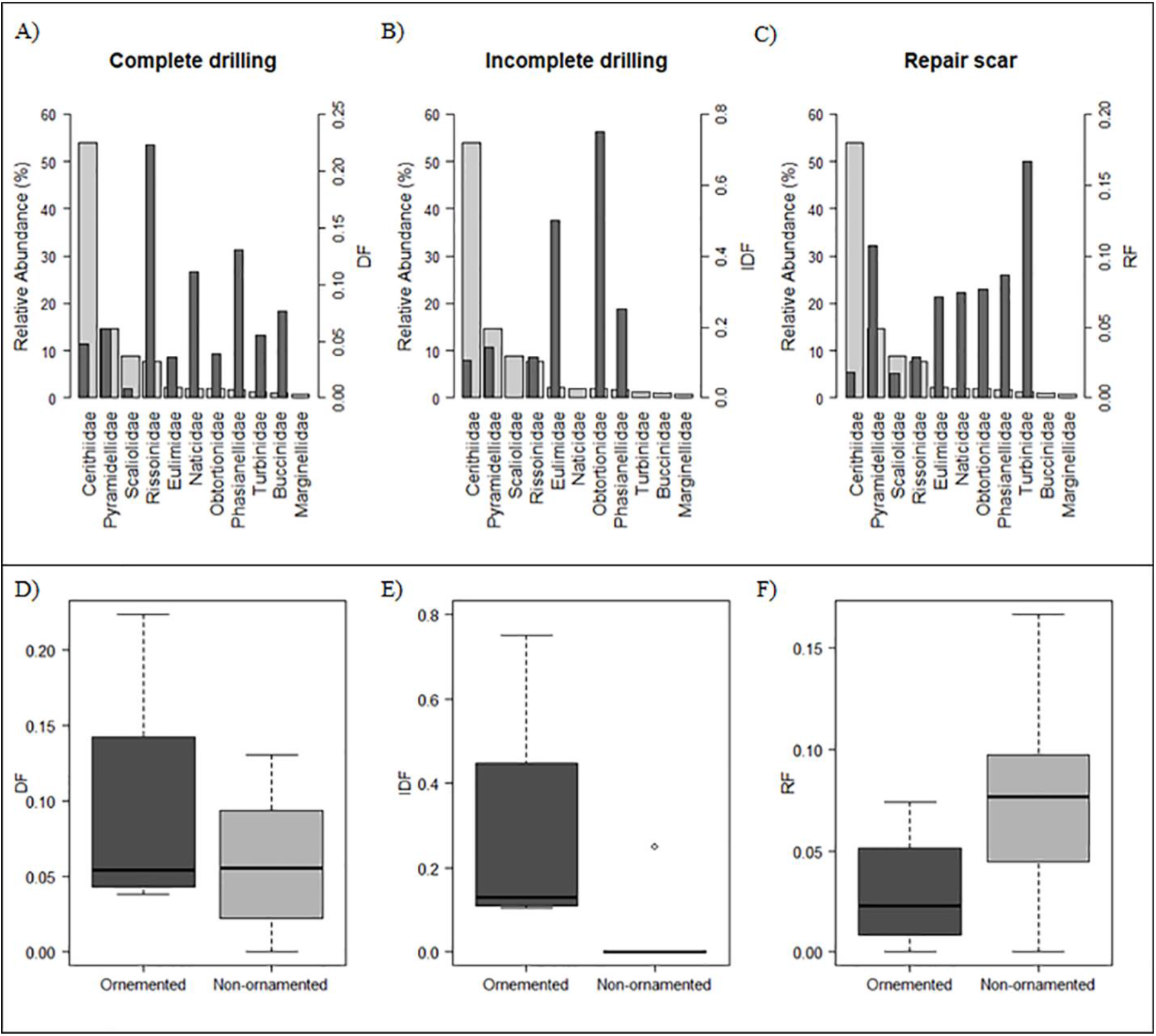
The variation of predation marks with the relative abundance of family (A-C) and nature of ornamentation (D-F). The frequency of complete drillhole, incomplete drillhole and repair scars are represented by panels from the left to right. The boxes in the bottom panel are defined by 25th and 75th quantiles; thick line represents the median value.

**TABLE 2.** Predation intensity in terms of drilling frequency, incomplete drilling frequency and repair frequency with respect to size.

| Size class | Drilling frequency (DF) | Incomplete drilling frequency (IDF) | Repair frequency (RF) |
| --- | --- | --- | --- |
| Small (< 1mm) | 0.05 | 0.07 | 0.02 |
| Medium (1-2mm) | 0.06 | 0.15 | 0.04 |
| Large (>2mm) | 0.09 | 0.32 | 0.11 |

**FIGURE 5.**
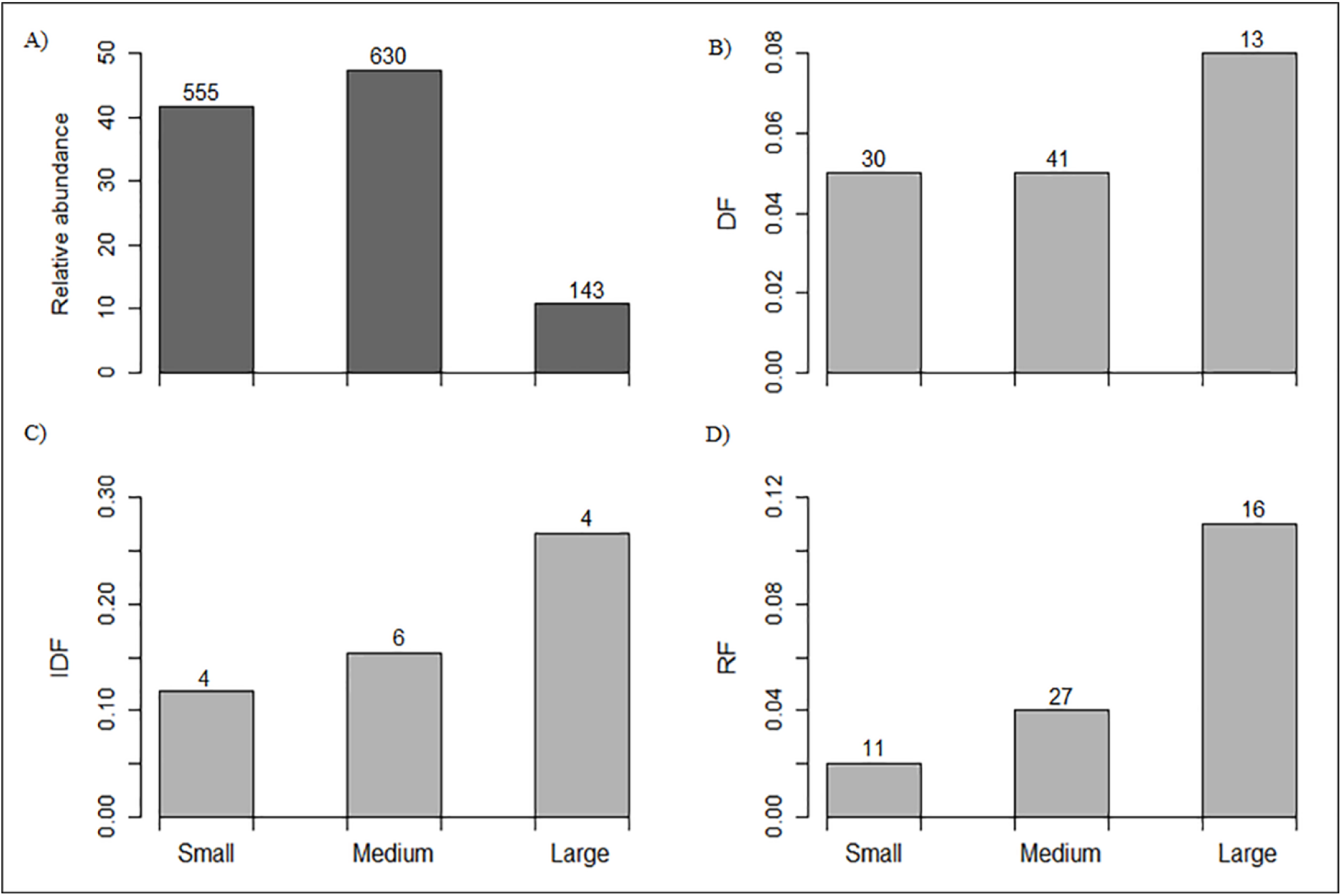
Histogram showing the size class distribution of A) drilled and undrilled specimens, B) drilling frequency (DF), C) incomplete drilling frequency (IDF), and D) repair frequency (RF).

**TABLE 3.** The results of the chi square tests done to evaluate the significance of variation in predation intensity interns of complete drilling incomplete drilling and repaired (significant results are marked in bold).

| Size class | Chi square value for DF | P-value for DF | Chi square value for IDF | P-value for IDF | Chi square value for RF | P-value for RF |
| --- | --- | --- | --- | --- | --- | --- |
| Small-Medium | 0.0086 | 0.9263 | 0.1961 | 0.65792 | 5.041 | **0.0247** |
| Medium-Large | 3.0754 | 0.079484 | 4.6882 | **0.03037** | 10.5719 | **0.00118** |
| Small-Large | 2.7708 | 0.096001 | 6.1884 | **0.01286** | 25.919 | **0.00001** |

**FIGURE 6.**
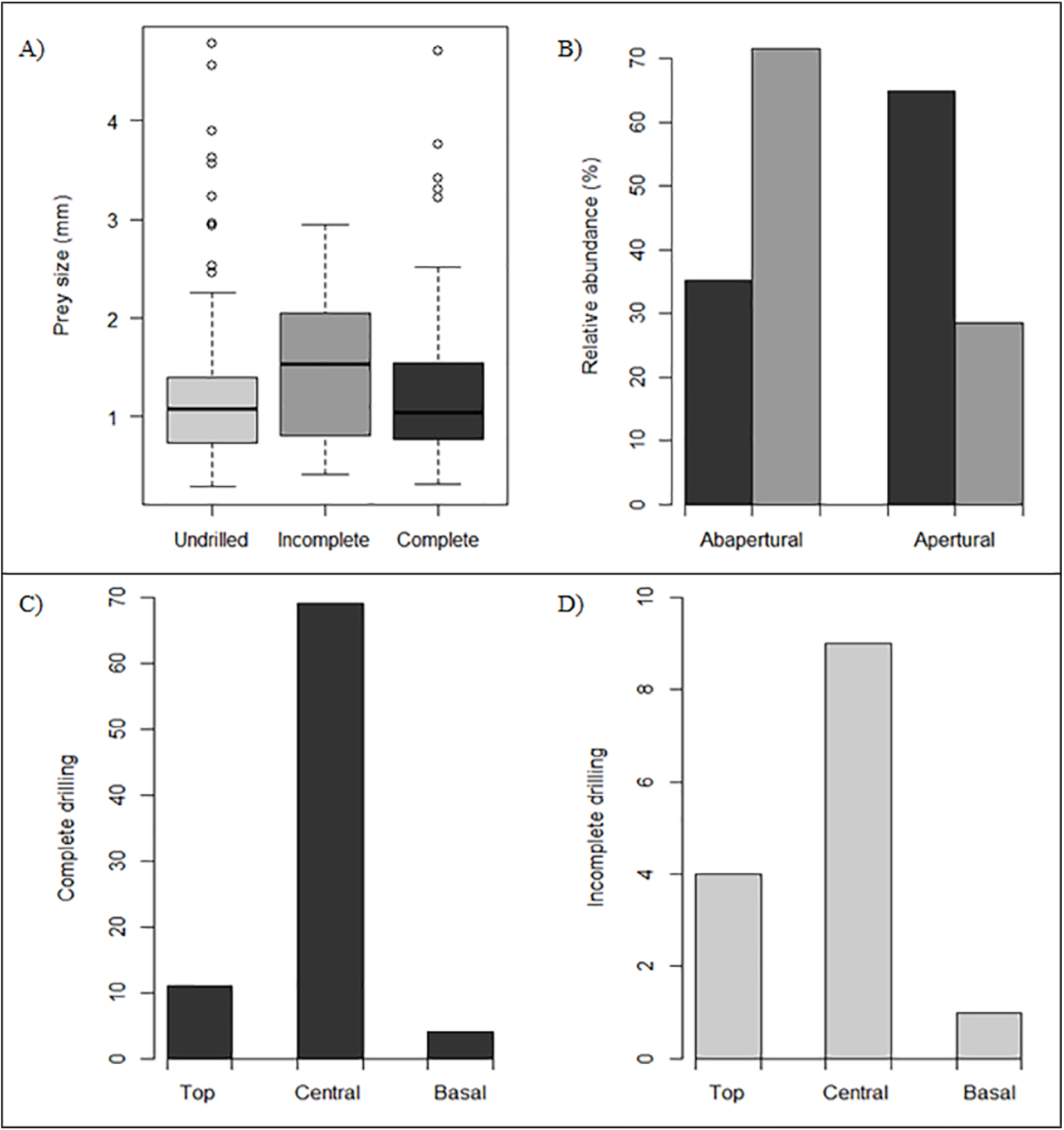
Plot representing the variation in A) prey size and B) site selection between successful and unsuccessful predation attempts. The boxes in A) are defined by 25th and 75th quantiles; thick line represents the median value. The bar plots in B) represent relative abundance of specimens with complete drillhole (black) and incomplete drill holes (grey).

**FIGURE 7.**
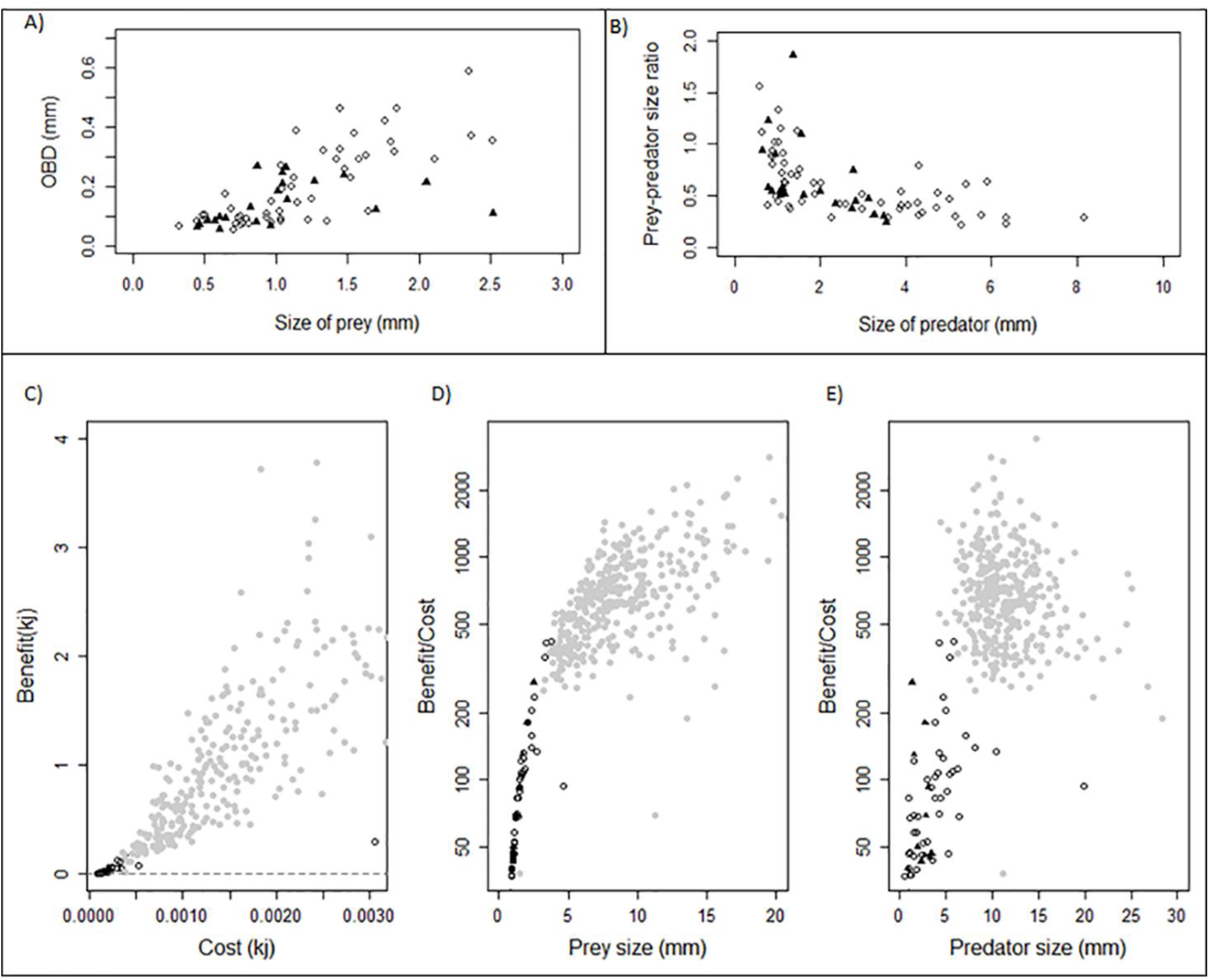
Plot showing the relationship between A) size of the prey and the outer borehole diameter, B) the size of the predator and the prey-predator size ratio. Open circles represent naticid drilling and the closed triangles represent the muricid drilling. Cost – benefit relation for the micro gastropods, C) indicates the benefit gained by the predator species for a particular cost, D) scatter plot representing the relation between prey size and the benefit / cost ratio, E) relation between the inferred predator size and the benefit / cost ratio. The dotted lines in the bottom panel indicate the minimum requirement for a successful predation (benefit = cost).

When compared to the other drilling predation observed in macrogastropods of Miocene (Table 4), DF of Quilon Limestone assemblage is lower compared to the other locations, except for Kutch (Goswami et al., 2020) (Table 4, Figure 8). The benefit-cost ratio is significantly higher for the macro gastropods of Kutch than the micro gastropods from Kerala (Wilcox test, p<<0.01) (Figure 7D, E). The family-level global comparison also shows a low DF for microgastropods in contrast to macrogastropods, except for the Rissoinidae family (Figure 9 A). Family-level comparison of RF demonstrates similar low-frequency in microgastropods (Figure 9 B).

**TABLE 4.**
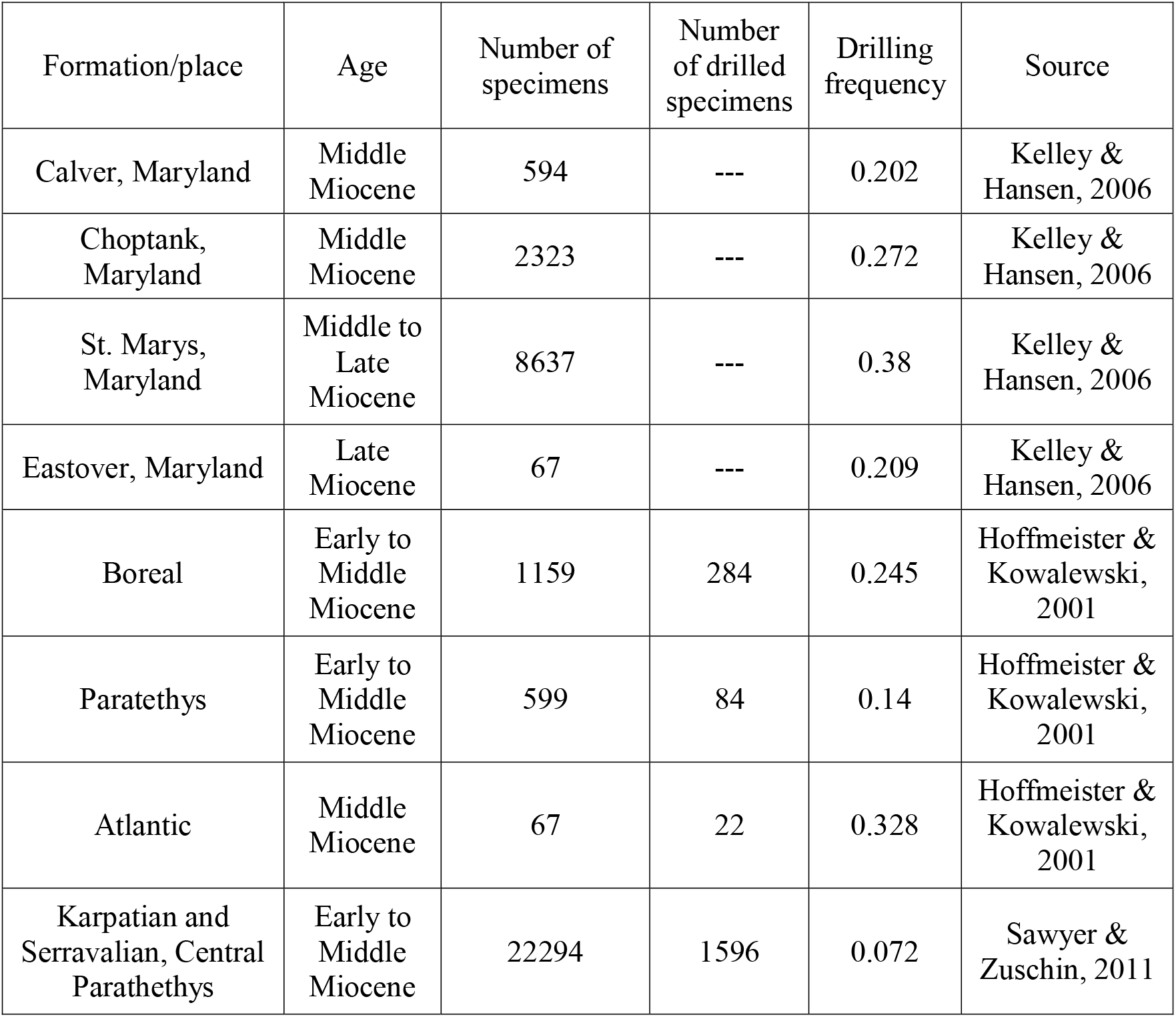
Spatiotemporal comparison of drilling predation data on gastropods from other major Miocene assemblages.

**FIGURE 8.**
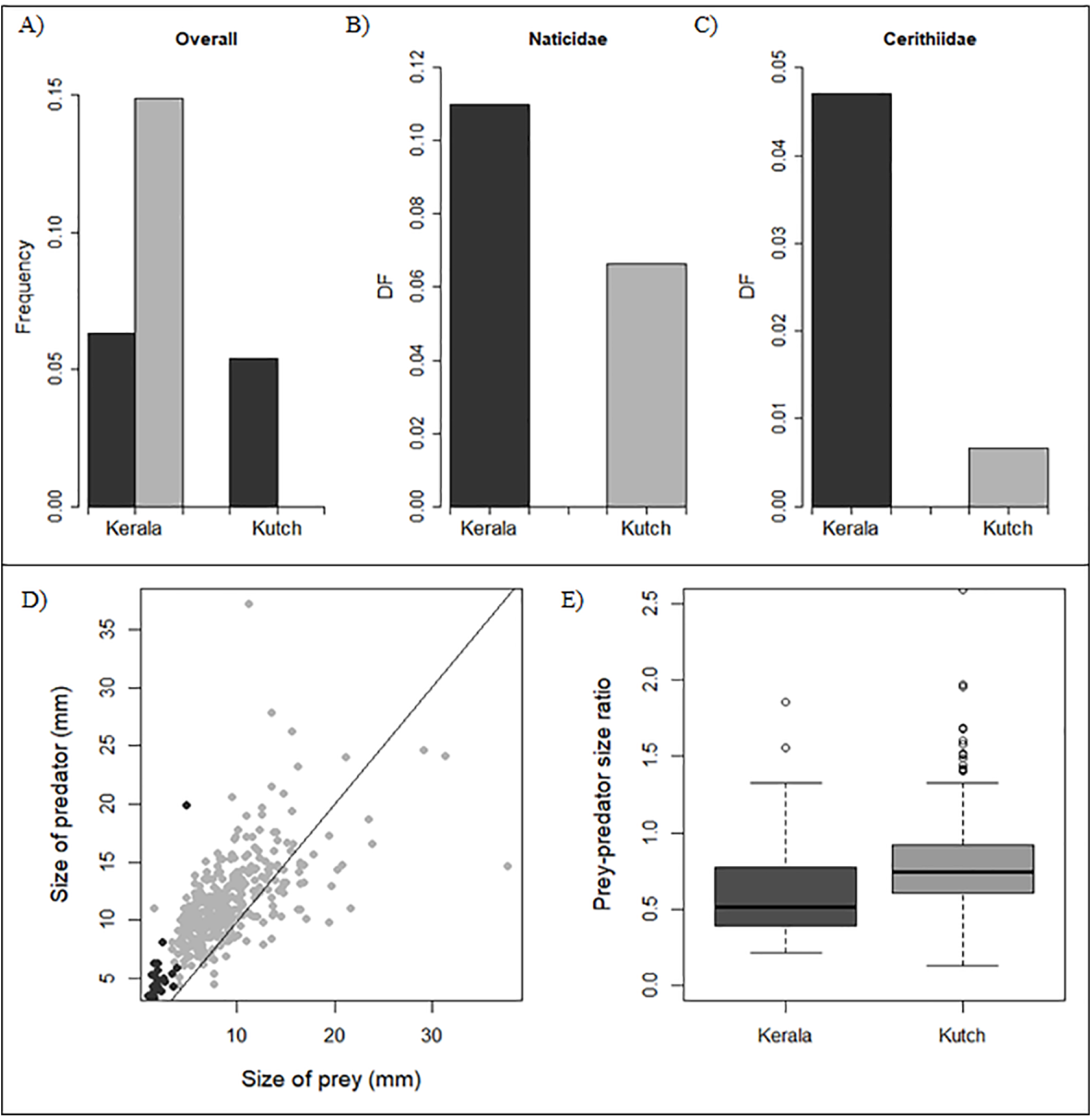
Predatory patterns in Kerala and Kutch A) comparison of DF and IDF, variation in DF for the common families B) Naticidae and C) Cerithiidae D) Relation between inferred size of the predator and size of the prey, grey represents Kerala specimens and black represents Kutch, E) boxplot representing the variation of prey predator size ratios.

**FIGURE 9.**
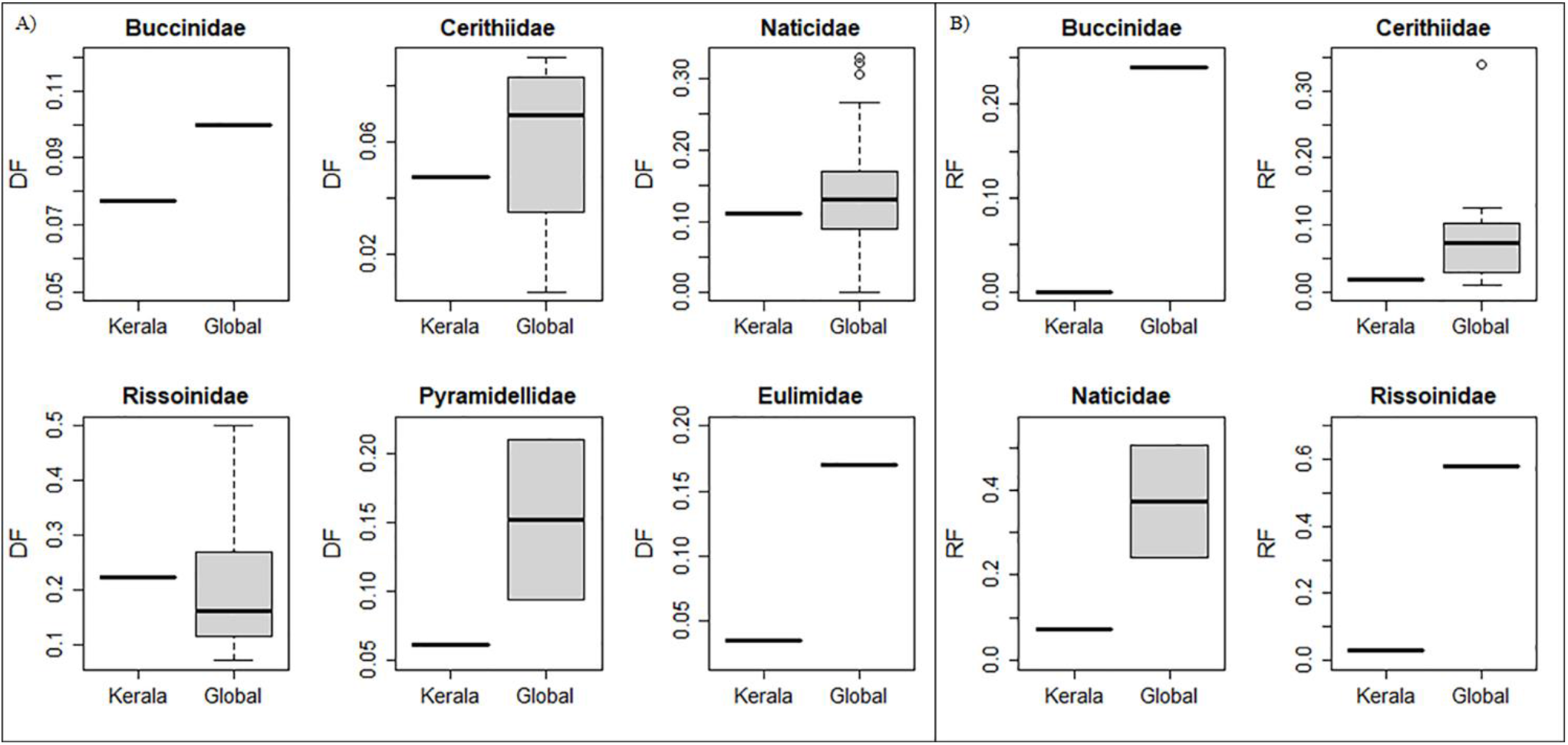
Family-specific comparison in predatory patterns between Kerala and the other coeval formations worldwide for A) DF and B) RF.

## DISCUSSION

Drilling predation on molluscan prey is the most common fossil record of predation followed by repair scars (Klompmaker et al. 2019). Temporal and spatial pattern of predation patterns has been established for molluscan species (Kelley and Hansen, 1993; Kelley and Hansen, 2006; Klompmaker et al., 2017) using a variety of approaches including controlled experiment (Chattopadhyay and Baumiller 2007; Chattopadhyay et al, 2014a, Das et al., 2015), ecological survey (Mondal et al., 2014; Pahari et al., 2016, Chattopadhyay et al, 2014b, 2015) along with documentation of fossil ecosystems. Despite such a large breadth of research on molluscan predation, micromolluscs are largely ignored. Individuals of ostracods and foraminifera that are comparable to micromollusc in size are known to be preyed upon drilling gastropods (Culver and Lipps, 2003; Reyment et al., 1987; Reyment and Elewa, 2003). It is, therefore, expected that micromolluscs will also be targeted by predators. Chattopadhyay et al (2020) documented the drilling predation on microbivalve prey from Quilon limestone and demonstrated the selective nature of drilling predation even in the micromolluscs for the first time. Although there have been studies on the evolution (Weigand et al., 2013) and habitat preferences (Olabarria et al., 2002) of microgastropods, there has not been any study on the predation patterns in microgastropods. The present study attempted to fill this gap.

### Predator identity

Naticid predators are responsible for the majority (76.5%) of the drill holes in the Quilon microgastropods as affirmed by the parabolic shape (Kabat 1990). The presence of individuals of the naticid family in our sample and the reported presence of multiple naticid genera (*Tanea, Natica*) in the assemblage (Harzhauzer 2014) confirm the identity of the naticid predators. Some of the naticids from the assemblage were as small as 0.035 mm, implying a shell size of approximately 0.4 mm; these could be juvenile naticids because the shells were extremely thin and lack any strong mineralization. The non-naticid drill holes had a straight cylindrical boundary indicating muricid predation (Carriker, 1981; Hoffman et al., 1974; Kabat, 1990). Although we did not find any muricid specimens in our sample, the presence of muricid family (*Triplex* and *Dermomurex*) in the same locality (Harzhauzer 2014) reveals the identity of the muricid drilling predator.

Repair scars are primarily produced after a non-lethal breakage often due to a failed predation attempt by fishes and crabs. The presence of *Xanthid* crabs in our specimens and reported presence from the same locality (Verma 1977) points towards a potential durophagous predator. The lack of presence of multiple repair marks and the higher number of repair marks among the ‘large’ microgastropods indicates a non-random predatory attack.

### Factors guiding the prey choice

The relative abundance of prey species is a good representation of the encounter frequency, and studies have suggested that the predation intensity may be linked to the prey availability (Leighton 2002, 2003). However, taxon-specific DF, IDF, and RF in our study are not correlated to its relative abundance (Figure 4 (A-C)) – a pattern consistent with findings for macro molluscs, both in the past and present ecosystems (Beu and Maxwell, 1990; Kelley et al, 2003; Mallick et al., 2014; Pahari et al., 2016). A lack of correlation between predation intensity and relative abundance indicates a predator’s preference towards a particular prey species, even if it is not the most abundant; such prey is often preferred by the predator due to certain morphological traits and highlights a selective behavior demonstrated by the predator. Our specimens show a highly selective nature of prey choice for both drilling and durophagous predation primarily guided by the morphological characters of the prey including size and ornamentation, for both drilling and durophagous predation.

#### Size

The size of an individual often dictates if it is targeted by a particular predator and determines the outcome of a predatory interaction. The reliable reconstruction of predator size is possible for drilling predation where experimental studies confirmed a strong positive correlation between the size of OBD and the size of the predator for specific families (Carriker and Gruber, 1999; Kitchell et al., 1981; Klompmaker et al., 2017; Kowalewski, 2004). The validity of the relationship has never been demonstrated for microgastropods. The inferred sizes of naticid and muricid predators in the microgastropod assemblage are comparable to the size of corresponding specimens found from the locality pointing to the validity of the approach.

Preys larger than a specific size are often avoided by predators due to difficulty in handling (Vermeij 1987). The smaller prey is thought to offer low energetic gain and hence, not selected. Consequently, the predator targets the medium-sized prey to maximize their energy gain (Boggs et al., 1984; Kelley, 1988; Kitchell et al., 1981; Chattopadhyay et al, 2020; Kitchell et al., 1981; Pahari et al., 2014). The low DF in the smallest size class in our sample supports this and suggests that the smaller size class is less likely to be attacked and it provides refuge from predation. However, the higher IDF and RF in the larger size class suggest that the larger prey is efficient in escaping the predator once it is attacked. This suggests a much complex prey-predator dynamics where smaller size class is not preferred and larger preys are more successful in escaping from predators. Among two families of drilling predators, the naticids show a significant positive correlation between individual predator size with the prey size demonstrating a strongly size-selective behavior (Figure 7 A, B). The absence of such size-selectivity of muricid predation is not unique to micro gastropods and has been reported from the macro gastropods (Tull and Böhning-Gaese, 1993).

#### Ornamentation

Surface ornamentation plays an important role in determining the outcome of a predatory encounter. The ornamentation increases the effective thickness of the shell, making it more difficult to drill through. The presence of surface ornamentation such as concentric ribs is found to reduce the incidence of successful drilling in bivalves (Klompmaker and Kelley 2015). Although insignificant for DF and RF, the IDF is found to be significantly higher in microgastropods with ornamentation suggesting that ornamentation increases the probability of drilling failure. The two non-ornamented families (Eulimidae and Phasainallidae) with higher IDF have a smooth shiny surface that is hard to grab. Moreover, Eulimidae are often associated with echinoderms that protect them from predators (Waren, 1983). A slightly higher RF (although statistically insignificant) was found among the non-ornamented specimens supporting the effect of ornamentation producing failures in durophagous attacks. The highest RF, however, is found in a non-ornamented microgastropod family - the Turbinidae. The small size and the smooth shell may have helped them to escape from the durophagous attacks.

#### Taxon

Both drilling and durophagous predators are known to demonstrate taxon selectivity (Chattopadhyay and Dutta, 2013, Chattopadhyay et al, 2015, Alexander and Dietl, 2003). Our study suggests that some prey taxa are preferred and the preference cannot be completely explained by the lack of morphological defense, such as the Rissoinidae family. They also have ornamentation such as ribs increasing their effective thickness, which should have acted against the predatory attacks. The abundance does not explain such higher rates always; families such as Scaliolidae and Cerithiidae have a larger population yet have a lower DF. In absence of obvious high encounter frequency or morphological weakness, their behavioral traits may have contributed to such increased predation pressure.

Individual predatory families also show distinct selective patterns. Muricids are found to prey heavily upon Obtorionidae. Kitchell et al (1981) have found that muricids are capable of drilling deeper holes enabling them to prey on molluscs with a thicker shell or higher ornamentation like the Obtortionidae. Because the deeper drill holes require longer drilling time, the probability of interruption by other predators and prey escape leading to incomplete drillings. This is also supported by the high IDF observed among Obtortionidae (Figure 3C, Figure 4B). In contrast to the overall dominance of naticid drilling, the assemblage demonstrates a low incidence of naticid cannibalistic behavior. Out of 64 naticid drillings, only two are cannibalistic and both of them are found in the small size class of prey supporting the experimental findings of the higher rate of cannibalism among smaller prey (Chattopadhyay et al, 2014).

### Predatory preference for site selection

Naticid predators often show stereotypic behavior in selecting the drilling site (Dietl and Alexander, 2005). The majority of the complete naticid drill holes are located in the central region of the microgastropods (48.6%) (Figure 6 C). Similar stereotypic behavior is known from macromolluscs (Allmon et al., 1990; Hagadorn and Boyajian, 1997; Goswami et al, 2020). When a prey species is alarmed, it withdraws the soft part inside the shell, up to nearly its central region (Hansen and Kelley, 1995; Kitchell, 1986). The drill holes in the central region ensure access to the soft tissue. A similar pattern is present among muricid drill holes suggesting a stereotypical behavior even of the muricid predators.

Our results show that drill holes are concentrated on the apertural side, mostly between the first and the fourth quadrant. Sometimes when site-selective behavior is observed, the second and third quadrant is mostly preferred. Such patterns are found since the prey species are on their dorsal side most of the time. The position of the drill hole is also dependent on the size and the morphology of the prey and the predator (Ansell, 1960; Kabat, 1990; Negus, 1975; Sohl, 1969). During naticid predation, the predator completely covers the prey to restrict its movement and it is often seen that they release a chemical that numbs the prey (Carriker and Gruber, 1999). Dietl and Alexander (2000) have explained that in confamilial predation in naticids they observe a significant number of drilling near the umbilicus because it would help the predator to immobilize a relatively “dangerous” prey, by covering the aperture using the foot. This could be observed even when the prey is significantly larger and mobile. The higher intensity of naticid drill holes on the ventral side of the shell in our data thus suggests a stereotypical behavior by the predator to effectively immobilize the prey.

### Prey effectiveness and repair frequency

The presence of incomplete drill holes, multiple drill holes, and repair-marks demonstrate the prey’s ability to escape, the inability of the predators to complete an attack due to an interruption (Kelley et al, 2003). Incomplete drill holes do not always indicate prey’s escape, because there are cases that reported the suffocation of the prey thus resulting in death (Hutchings and Herbert, 2013). The results indicate a significant increase in IDF and RF with size. This may suggest that the shell thickness of the larger prey might be slightly higher, making it less desirable. So these higher rates represent prey’s physical defense mechanism acquired over its lifetime to escape predation. Chattopadhyay and Baumiller (2007) showed that the presence of secondary predators may result in the abandonment of the prey by the predatory gastropods, leading to the development of incomplete drilling. In such cases, RF is proportional to IDF and inversely proportional to DF (Chattopadhyay and Baumiller, 2010). The microgastropod assemblage, however, does not show any significant correlation (p = 0.43 for DF-RF, and p = 0.92 for IDF-RF) between these three indices suggesting limited involvement of predatory abandonment in producing incomplete drill hole.

A high RF of an assemblage indicates the presence of a stronger predator (Vermeij et al., 1981). We have standardized the RF for both size and taxon. For the size standardized calculation, two reasons could account for the higher RF in the large size class: a) larger preys are usually the older and hence, accumulate the scars developed over multiple attacks during its ontogeny, b) larger prey are more likely to survive an attack in comparison to smaller prey and hence carry the signature of non-lethal attack. Multiple drill holes and incomplete drilling are not uncommon in the assemblage. Lower IDF and MULT values from macrogastropod assemblages of Miocene have been interpreted as the signature of highly efficient predation (Fortunato 2007). The relatively higher values of IDF (14.3%) and MULT (9.7%), compared to the Kutch assemblage (0% and 0.70%) (Goswami et al 20219) along with lower DF indicate that micro gastropods have an effective way of escaping predation.

### Energetics of the predation

The non-random prey selection by predation is explained by the cost-benefit principle (DeAngelis and Kitchell, 1985). The cost is the invested energy by the predator in finding, capturing, and consuming the prey; the benefit is the energetic value of the prey tissue by the predator. The principle suggests that a predator selects prey to maximize the net energy gain i.e. the difference between the benefit and cost. This principle has been shown to operate in the prey selection by both naticids (Kitchell et al., 1981) and muricids (Chattopadhyay and Baumiller, 2009) on macromolluscan prey. Cost-benefit analyses confirm that the selection in micromolluscan prey is non-random and each of the successful predations yielded a positive net energy gain (Figure 7A). The microgastropod prey yield higher energetic gain with increasing size primarily because of the increase in soft tissue volume and a negligible increase in thickness of the prey (Fig A, D). This results in the exponential increase in benefit: cost ratio with prey size. This explains why smaller sizes among microgastropods are not the preferred prey confirmed by the lower DF in comparison to larger size classes (Figure 7(C-E)). It is also important to note that none of the individuals below 0.35mm are drilled. This also confirmed the “negative size refuge” exists in microgastropods similar to microbivalve prey (Chattopadhyay et al, 2020). The cost-benefit analysis also confirms that the micromorphy may act as an effective defense strategy by making the smaller sizes less preferred.

The cost-benefit analysis also brings out interesting behavioral attributes of the predator. Although the prey-predator size ratio decreases with predator size (Figure 7B), the net energy gain increases. This implies that smaller predators, despite their selection of relatively larger prey, do not benefit energetically due to a disproportionately higher metabolic cost. When compared between two families of drillers, naticid drillings are more beneficial than muricids; the naticids are found to have a significantly higher net energy gain compared to muricids.

### A comparison to macro gastropods

Low values of drilling frequency in microbivalves in comparison to coeval global averages have been used to establish the effectiveness of micromorphy against drilling predation (Chattopadhyay et al, 2020). Our study confirms this finding for both drilling and durophagous predation. The low predation intensity in family-level comparison with macrogastropods indicates the predation resistance of microgastropods (Figure 9). Such lower intensity among the microgastropods is probably driven by the low energetic yield making them less preferred as demonstrated by the cost-benefit analyses. This is also supported by the higher benefit-cost ratio, observed among the macro gastropods from Kutch (Figure 7 C-E). However, there might be other factors that could affect the intensity of predation.

The studied section is interpreted to represent a seagrass environment (Reuter et al., 2011). Seagrass environment is often found to provide a natural refuge from the predators (Irlandi 1997; Wall et al., 2008) where leaf blades diminish the mobility and visibility of the predators (Heck and Thoman, 1981; Irlandi, 1997). The roots also prevent the digging and hence, limiting the activity of infaunal predators (Wall et al., 2008). Since many of the predators (muricid, xanthid crab) are epifaunal, the effect of the seagrass cannot completely explain the low predation intensity of Quilon microgastropod assemblage.

Differential preservation of the macro-and microgastropods may also contribute to the observed low predation intensity of microgastropods. Generally, the smaller gastropods, especially juveniles, are rarely preserved in fossil records (Cooper et al., 2006; Kidwell, 2001) often leading to a difference in observed predation intensity across size classes (Chattopadhyay et al, 2016). One of the taphonomic biases known to lower the inferred DF is the differential shell strength of drilled and undrilled shells. Drill holes reduce the shell strength and make the drilled shells more susceptible to compression-induced breakage potentially leading to a reduced DF (Roy et al., 1994). However, the difference in breaking load between drilled and undrilled shells is more pronounced in larger shells (Roy et al., 1994, Fig. 3) – a pattern that is more likely to lower DF in macromolluscs. Moreover, the lighter shells of microgastropods are likely to be carried as suspension load in contrast to the microgastropods that travel as bed load and get reworked in the process (Reuter et al., 2011). Most microgastropods in our sample or reported collection retained their original structure without any breakage pointing to the limited role of compaction-induced breakage in developing the assemblage. Apart from this, the difference in hydrodynamic properties of drilled and undrilled shells are also known to create assemblages with reduced DF (Chattopadhyay et al, 2013a, b). However, the difference is more pronounced for larger size classes (Chattopadhyay et al, 2013b, Fig. 5). Both the taphonomic attributes (compaction, hydrodynamics) that are known to reduce DF are more likely to affect microgastropods and do not explain the observed low predation intensity in microgastropods implying a relatively negligible role of taphonomy in creating the pattern.

The relative abundance of predatory species is known to explain the predation intensity of a region (Allmon et al., 1990; Kardon 1998; Sawyer and Zuschin, 2011). In the recent study by Goswami et al. (2020), the low drilling intensity of macrogastropods from Kutch is explained by the low abundance of predators. Because of the low abundance of muricid gastropod in their assemblage, most muricid-like drill holes have been attributed to naticid identity. Microgastropod assemblage of Quilon limestone is characterized by a lower relative abundance of potential drillers (2.04%) in comparison to the reported values from other Miocene assemblages, such as Kutch (2.27-4.55%) (Goswami et al., 2020). Muricid drilling is present in our collection and muricid specimens have been reported from the same locality (Harzhauser, 2014). However, the absence of muricid gastropod specimen in our documented collection is a probable indicator of its lower abundance.

Apart from the relative abundance of predators, the absence of preferred prey may also result in low predation intensity. The Quilon assemblage reports a fewer number of Turritellids – a species known to be a preferred prey with high DF (Goswami et al 2020, Fortunato, 2007; Kojumdjieva, 1974) thereby making it a preferred prey. The absence of this group may have contributed to the overall lower DF of the Quilon assemblage. The availability of other preferred prey may also contribute to the lower predation intensity among microgastropods. Chattopadhyay et al (2020) have reported the drilling predation among the micro bivalves from the same locality with similar predation intensity (DF=0.06) making it unlikely to be a preferred prey over microgastropods. Other potential preys of this size class include ostracods and forams. The thin shells of the ostracods might lower the energy for drilling making them desirable prey (Culver and Lipps, 2003; Reyment and Elewa, 2003; Reyment et al., 1987). Although we do not have any direct evidence of predation from these groups, the high abundance of ostracods (Yasuhara et al, 2020) and foraminifera (Rögl and Briguglio, 2018; Briguglio and Rögl, 2018) have been reported from the Quilon assemblage and supports the availability of alternate prey types. This also opens the possibility for future studies to explore the predatory interaction in these groups to understand predator-prey dynamics at extremely small size classes.

## CONCLUSION

Predation in the molluscan communities, from the recent and the past ecosystem, is studied in-depth, with the possible exception of micro molluscs. The present study attempted to fill this gap by studying the predation signature in micro gastropods from the Quilon limestone bed of Kerala. The predation intensity of this assemblage is quite low for drilling (DF= 0.06) and durophagous (RF=0.04) predation. Also, the repair frequency (RF) and the incomplete drilling frequency (IDF) are found to be lower for the micro gastropods in comparison to family-specific values of global reports. These results support the previous findings of micromorphy acting against drilling predators with low drilling predation intensity as shown among micro bivalves (Chattopadhyay et al, 2020). The small size of the prey species is a good defense against predation, and inverse size refugia are observed among micro gastropods. However, the larger prey is found to escape the predation more efficiently as demonstrated by a higher IDF among large size class. The physical features of the gastropods affect the intensity of predation rather than the abundance. The lower intensity of predation in this size range might be a result of multiple factors that includes the lower number of predators, the seagrass environment, and the presence of other prey species. Finally, the cost-benefit analysis suggests an increasing benefit to cost ratio with increasing prey size explaining the potential reason for preferring microgastropods over microgastropods leading to the low predation intensity observed among micromolluscs.

## ACKNOWLEDGEMENTS

This work was supported by the Academic Research Grant of IISER Kolkata (ARF 2018-19), and DST Inspire fellowship. We thank Debarati Chattopdhyay and Venugopal S Kella for collecting the samples and initial processing.

## Notes

### Competing Interest Statement

The authors have declared no competing interest.

